# Primary metabolism determines the outcome of salicylic acid-mediated immune induction

**DOI:** 10.1101/2025.10.13.682132

**Authors:** Qian Zhang, Yucong Xie, Sargis Karapetyan, Jinlong Wang, Musoki Mwimba, Heejin Yoo, Xinnian Dong

## Abstract

Controlling the deleterious effects of immune responses is as vital as fighting infection. In plants, this is achieved, in part, by circadian clock-mediated regulation, such as the synthesis of and response to the immune hormone salicylic acid (SA)^1,2^. Application of SA at the same concentration under light/dark cycles induces immunity with minimal impact on growth, however, prolonged darkness leads to plant death^2^. To uncover what determines this life-or-death outcome, we identified twenty *survival of SA-induced death* (*ssd*) mutants through genetic screening. These mutants are defective in starch, glucose, and nitrate metabolism, and circadian regulation, and accumulate excessive starch and/or glucose. Likewise, glucose application rescues SA-treated plants in prolonged darkness. Surprisingly, SA treatment does not deplete glucose, but instead, induces amino acid and fatty acid catabolism. Through transcriptomic analyses of glucose-rescued WT plants and *ssd* mutants for shared pathways, we found that SA triggers plant death in darkness by inducing oxidative stress, and water loss, while glucose antagonizes these processes, boosts ER protein processing and re-establishes the anabolism-catabolism balance. Interestingly, the programmed cell death induced by effector-triggered immunity shares common transcriptomic patterns with those observed during SA-induced cell death in darkness and could also be attenuated by glucose treatment. Therefore, coordination with the cellular metabolic context plays a central role in determining immune outcomes and optimizing plant health.

## MAIN

In plants, salicylic acid (SA) synthesis is induced upon pathogen challenge to promote broad-spectrum systemic acquired resistance (SAR) through activation of the signalling protein NONEXPRESSOR OF PATHOGENESIS-RELATED GENES 1 (NPR1)^3,4^. Interestingly, in the absence of pathogen challenge, the basal SA synthesis is directly controlled by the circadian clock transcription factor (TF), CCA1 HIKING EXPEDITION (CHE), to peak before dawn^1^ while the plant response to external application of SA is gated by the clock towards the morning^2^. This suggests that the temporal regulation of SA responses might be a strategy to enhance defence when the threat of infection is the highest, while mitigating its detrimental effects on plant growth and development. Indeed, while application of SA to plants under light/dark (LD) conditions induces immune responses with only occasional growth retardation^5^, it triggers plant death in prolonged darkness (DD)^2^. This result suggests that carbohydrate deprivation in DD may exacerbate the deleterious effects of SA. It also indicates the presence of a mechanism that coordinates plant immune responses with the daily metabolic activities, such as carbohydrate oscillations. In the daytime, glucose produced by photosynthesis is polymerized to transitory starch for storage in chloroplasts. At night, transient starch is degraded to provide glucose to support growth and this turnover is controlled by the circadian clock to ensure that starch is exhausted precisely at dawn^6^. The availability of glucose serves dual roles both as an energy source for primary metabolism and as a signalling molecule that coordinates growth with environmental conditions. Sensing of glucose through the HEXOKINASE1 (HXK1) or the TARGET OF RAPAMYCIN (TOR) pathway promotes anabolic processes and plant growth, while energy deficiency activates SNF1-RELATED PROTEIN KINASE 1 (SnRK1), which inhibits the TOR function and activates autophagy and catabolic processes to provide alternative energy sources^7^. However, whether and how primary metabolism impacts the outcome of SA-mediated immune induction remains a mystery.

Through a forward genetic screen, we identified twenty survival of SA-induced death (*ssd*) mutants in DD, majority of which were deficient in transient starch turnover, accumulating excess starch and/or glucose. Furthermore, exogenous application of glucose rescued the wild-type (WT) plants from SA-induced death. These results demonstrate that carbohydrate supply attenuates the stress caused by SA and protect the plants from the deleterious effects accompanying SA-mediated immunity. Further transcriptomic and chemical intervention analyses demonstrate that SA induces oxidative stress, inhibits anabolism and promotes protein and amino acid catabolism, followed by water loss and eventually plant death, while glucose suppresses these responses and upregulates ER processing and translation. Interestingly, many of the transcriptomic changes during SA-induced death in DD mirror those induced during effector-triggered immunity (ETI), which leads to localized programmed cell death (PCD) as part of the plant’s defence mechanism. Glucose application can also reduce ETI-associated cell death, highlighting the integral role that cellular metabolism plays in fine-tuning immune responses for optimal plant health.

### Forward genetic screen identifies *ssd* mutants in five distinct biological pathways

To elucidate whether SA-induced death in DD is caused by defence activation, we first tested the *npr1* mutant which is deficient in SA-mediated immunity^3^, and found a 100% death rate, comparable to the WT plants (Fig. 1a,b), suggesting that SA-induced death under DD is independent of NPR1. Therefore, to identify the molecular mechanisms behind SA-induced death in darkness, we performed a forward genetic screen with 1 mM SA application for surviving mutants, using seeds collected from ∼ 4,000 individual M1 plants after ethylmethanesulfonate (EMS) mutagenesis in the WT *Arabidopsis* background (Extended Data Fig. 1). Our screen identified twenty independent *ssd* mutants named *ssd1* to *ssd20*, with death rates ranging from 0% to 85%, lower than 97.2% in WT. Bulk segregant genome sequencing^8^ identified causal mutations in genes functioning across five distinct pathways, with some genes found mutated in multiple *ssd* mutants: transitory starch degradation [*ssd1*-*ssd5* in *DISPROPORTIONATING ENZYME 2* (*DPE2*), *ssd6*-*ssd10* in *BETA-AMYLASE 4* (*BAM4*), *ssd11* and *ssd12* in *ISOAMYLASE 3* (*ISA3*), *ssd13* in *STARCH EXCESS 1* (*SEX1*), *ssd14* in *BETA-AMYLASE 3* (*BAM3*), *ssd15* in *LIKE SEX4 1* (*LSF1*)], cellulose biosynthesis [*ssd16* in *CELLULOSE SYNTHASE CATALYTIC SUBUNIT 7* (*CESA7*)], brassinosteroid (BR) signalling [*ssd17* in *BR INSENSITIVE 1* (*BRI1*), *ssd18* in *DWARF 5* (*DWF5*)], nitrate metabolism [*ssd19* in *NITRATE REDUCTASE 2* (*NIA2*)], and circadian regulation [*ssd20* in *EARLY FLOWERING 3* (*ELF3*)] (Fig. 1c,d and Supplementary table 1).

**Fig. 1.**
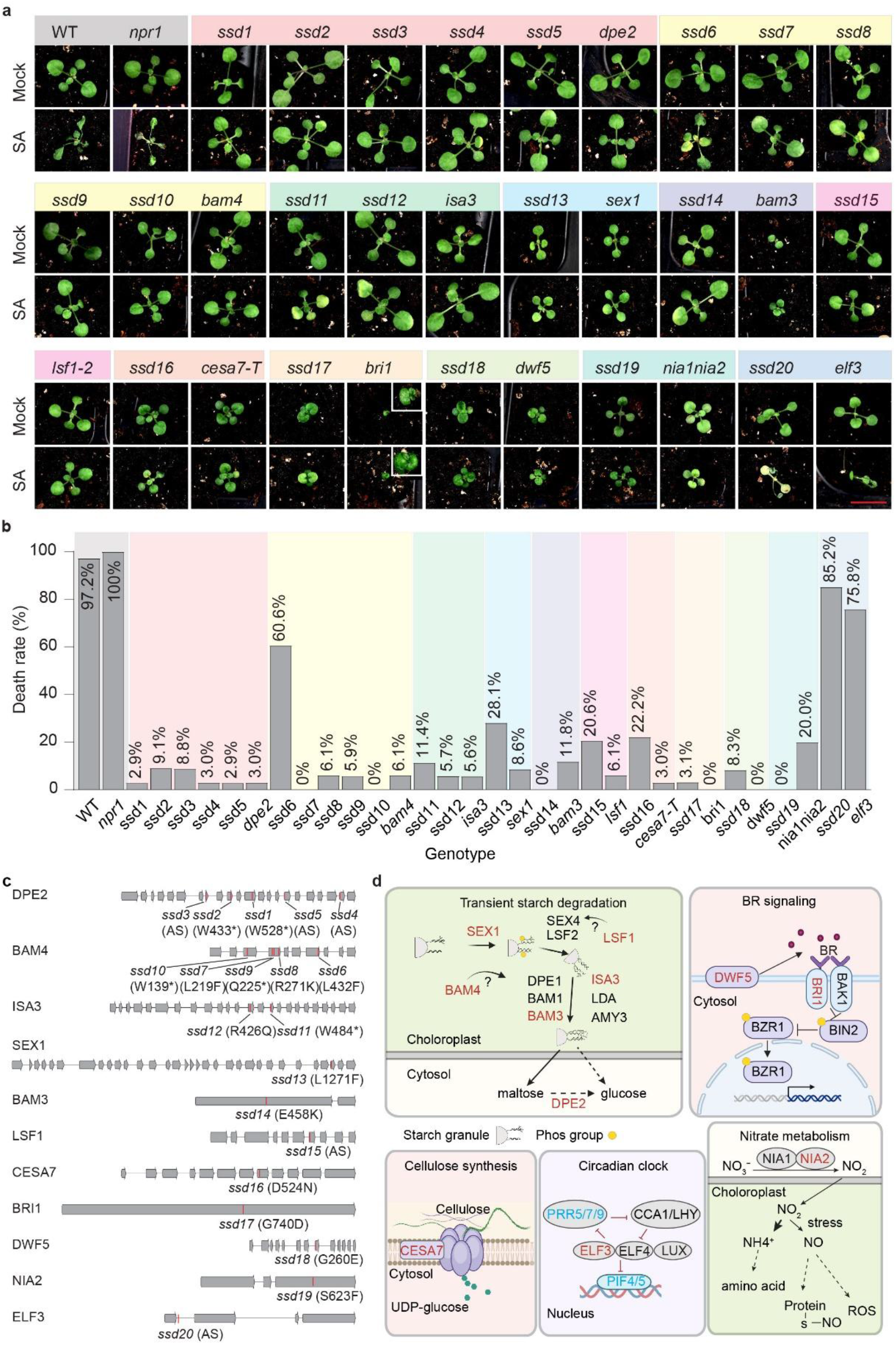
The *ssd* mutations rescue SA-induced death in prolonged darkness (DD) by disrupting distinct metabolic pathways. (a) Representative images of *ssd* mutants and their corresponding T-DNA insertion lines (with previously known gene names) after mock or 1 mM SA treatment. Plants were grown in soil in light/dark (LD) cycles for 14 days before SA treatment at the end of the dark cycle, and the plant phenotypes were scored 72 h after incubation in prolonged darkness. Scale bar = 1 cm. (b) Quantification of death rates from (a). Approximately 36 plants/genotype were used for quantification of the death rate in all figures. (c) Diagrams illustrating *ssd* mutations mapped onto their respective gene structures. Letters and numbers in parenthesis indicate the site and resulting amino acid changes for each corresponding *ssd* mutation. *, nonsense mutations; AS, mutations in alternative splicing sites. (d) Functional classification of all the *SSD* genes. Red letters represent genes identified in the *ssd* mutant screen and blue letters indicate additional genes tested through reverse genetics. The death rates for all the mutants presented in this and subsequent figures were assessed in at least three independent experiments with similar results.

To confirm the phenotype and exclude potential effects from secondary mutations in the *ssd* mutants, we examined the phenotypes of corresponding T-DNA mutants from the *Arabidopsis* mutant collection for all the *ssd* candidate genes. Except for *nia1/2* and *elf3* which showed death rates of 20% and 75.8%, respectively, other T-DNA mutants exhibited death rates close to 0% (Fig. 1b), confirming our genetic screen. The partial death phenotype in *nia1/2* and *elf3* mutants suggests possible functional redundancy in the nitrate metabolism and the circadian clock genes.

### Sugar application rescues SA-induced death

Mutations in the transitory starch-degradation enzymes have been known to result in excess starch accumulation^9^. As the majority of the *ssd* mutants are involved in starch degradation, we measured the starch amount by staining all *ssd* mutants and their corresponding T-DNA lines with the Lugol solution at dawn, when transitory starch is expected to be depleted in WT plants. In contrast to the WT, all starch degradation mutants (*dpe2, bam3, bam4, isa3, sex1*, and *isf1*) and the BR signalling *bri1* mutant retained detectable levels of starch in the morning, whereas *cesa7*, *dwf5*, *nia1/2,* and *elf3* mutants showed minor differences compared to WT (Fig. 2a and Extended Data Fig. 2a). Since the *ssd6-10* mutant alleles of *BAM4* showed variable starch staining, we performed a quantitative starch measurement in them and found significantly higher levels of starch in all alleles at both dawn and dusk (Fig. 2b), despite the weak staining results in *ssd6* (Extended Data Fig. 2a).

**Fig. 2.**
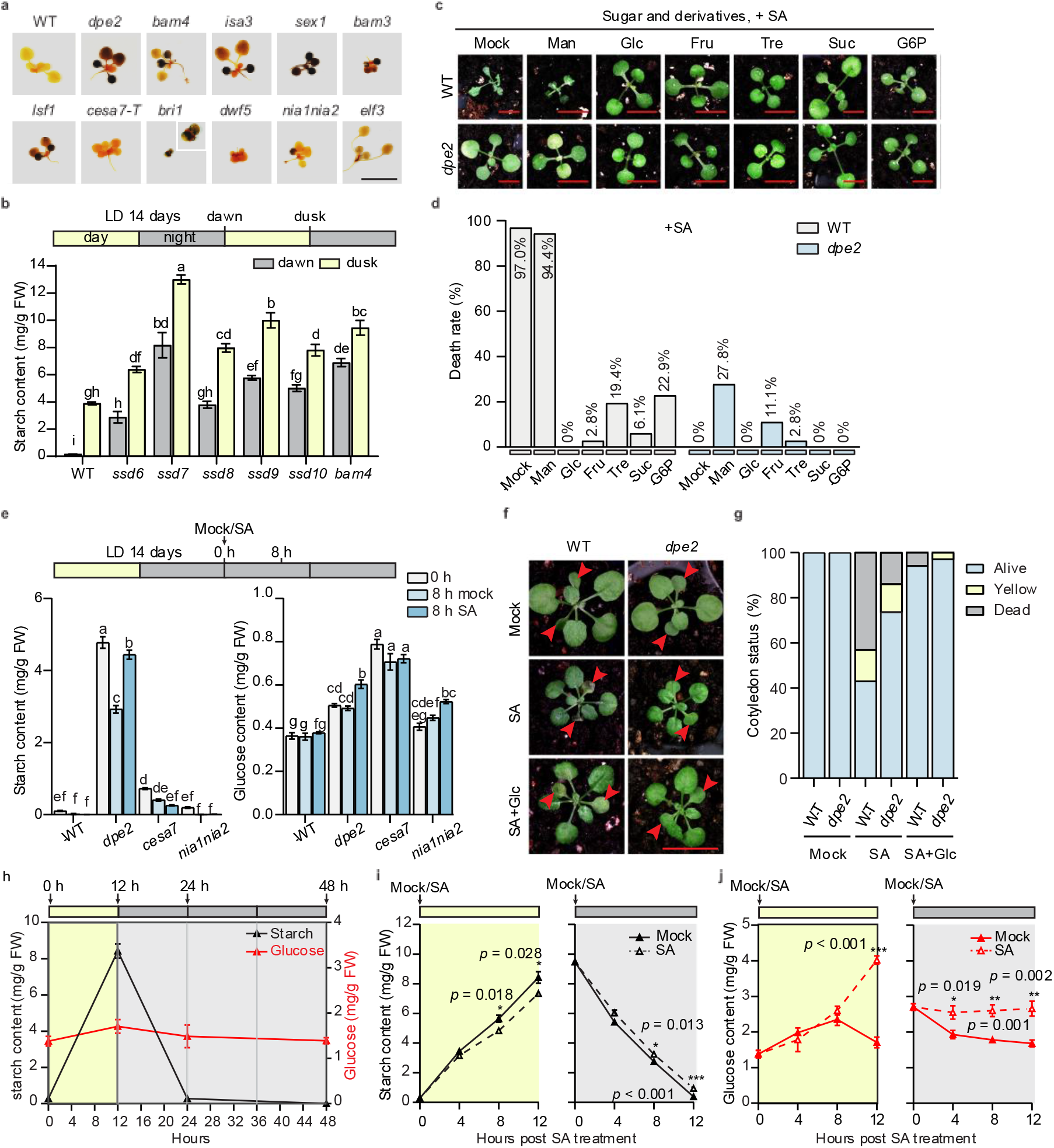
Glucose rescues SA-induced phenotypes in WT plants but SA does not deplete glucose. (a) Starch staining in the T-DNA mutants of the corresponding *SSD* genes. Plants grown in soil under LD cycles for 14 days were collected at dawn and stained using the Lugol reagent. Scale bar = 1 cm. (b) Starch quantification in *ssd6-10* and *BAM4* T-DNA mutant at dawn and dusk. On the top, the horizontal yellow (day) and grey (night) bars indicate times of the day for sample collection (or treatment for the following figures). (c) Exogenous application of glucose or related sugars/derivatives rescue plants from SA-induced death. Plants were grown in LD for 14 days and applied with 100 mM sugars/derivatives together with 1 mM of SA at the end of dark cycle and phenotypes were scored 72 h after incubation in DD. Mannitol (Man) was used at the same concentration as an osmotic control. Glc, glucose; Fru, fructose; Tre, trehalose; G6P, glucose-6-phosphate. Scale bar = 0.5 cm. (d) Quantification of death rates from (c). (e) Starch and glucose quantification in representative metabolic mutants with mock or SA treatment under DD. (f) Co-application of glucose rescues SA-induced phenotypes in the cotyledons when grown under LD conditions. Plants grown in LD for 14 days were repeatedly treated with mock, SA or SA + Glc for 7 days at the end of each dark cycle. Cotyledons were shown with red arrow heads. About 72 cotyledons from 36 plants were assessed for each treatment. (g) Quantification of dead, yellow and alive cotyledons from (f). (h) Starch and glucose dynamics during LD cycles and under DD. (i and j) Starch (i) and glucose (j) dynamics with mock and SA treatment at the start of day or night. All data are presented as mean ± s.e.m. Individual columns were compared using one-way ANOVA with Tukey’s post-hoc (b and e), different lowercase letters indicate statistical significance at p < 0.05; or using unpaired multiple two-tailed t-tests for comparing mock-and SA-treated samples (i and j), *p < 0.05, **p < 0.01, ***p<0.001. Six biological replicates were performed for each sample for starch and glucose assessment (b, e, h-j).

Based on these results, we hypothesized that starch overaccumulation in the *ssd* mutants possibly provides sugars or sugar signals that rescued them from death. To test this hypothesis, we sprayed plants with carbohydrates together with SA induction. We found that glucose, fructose, and sucrose dramatically reduced death rates in WT to 0%, 2.8%, and 6.1%, respectively, while trehalose provided a moderate level of protection (19.4% mortality).

Phosphorylated glucose (glucose-6-phosphate) also offered partial protection with a death rate of 22.9% (Fig. 2c,d). Mannitol, as an osmotic control, did not rescue WT (94.4% death rate), and even slightly decreased the survival of the *dpe2* mutant (27.8% death rate) (Fig. 2d). Moreover, even without SA, mannitol application caused low levels of cell death in both WT and *dpe2* in DD (Extended Data Fig. 2b,c). These results ruled out high osmotic pressure generated by the sugar solution as a direct factor for plant survival. Instead, they support our hypothesis that a supply of carbohydrate is a vital factor in rescuing SA-induced death in DD. Consistently, we found that compared to WT, the *dpe2* mutant accumulated both excess starch and glucose, while *cesa7* and *nia1/2* mutants, which did not accumulate extra starch, had higher glucose levels with SA treatment (Fig. 2e).

To determine whether glucose rescue is specific to SA-induced plant death in DD, we tested plants grown under LD conditions with repeated SA spray at the end of each dark cycle, when transitory starch is exhausted. We observed that cotyledons began to yellow and die after 7 sprays, while co-application with glucose rescued the phenotype (Fig. 2f,g). This indicates that that glucose mitigates SA-induced death under both DD and LD conditions. These findings also suggest that the plant death observed under DD is unlikely to be an emergent DD-specific phenotype, but rather an exacerbated side-effect of SA induction.

Since starch/glucose accumulation rescues plants from death after SA treatment in DD, we then tested whether this death was due to accelerated glucose consumption. We first measured the dynamics of starch and glucose in WT during the LD cycles and under DD. The results showed that starch accumulated during the day was depleted by the end of the night and remained undetectable under DD (Fig. 2h). In contrast, glucose levels, with only a slight increase during the day, remained relatively constant throughout the LD cycles and even after 24 hours of DD (Fig. 2h). Surprisingly, while SA application did not strongly affect starch turnover (Fig. 2i and Extended Data Fig. 2d), it significantly increased, rather than decreased, the glucose content in WT plants during both light and dark periods (Fig. 2j). Therefore, although extra glucose applied to WT or over-accumulated in the *ssd* mutants may contribute to plant survival, SA does not induce plant death through glucose and starch depletion.

### Glucose antagonizes SA-induced transcriptomic changes

To understand the underlying mechanisms determining this death-or-survival binary outcome from SA application alone or together with glucose, we performed quantitative RNA sequencing (QuantSeq) on WT plants treated with mock (M), SA (S), glucose (G), and SA plus glucose (SG) under both DD and LD, with sample collections at 8 hours and 24 hours post treatment (Fig. 3a). Principal component analysis (PCA) revealed that DD samples showed greater separation compared to LD samples, with SA and glucose treatments moving the overall transcriptome patterns in the opposite directions along PC1. Crucially, SG samples clustered closer to M and G than S samples under DD conditions (Fig. 3a), suggesting that glucose suppressed the effects of SA in DD. We then performed a Pearson correlation analysis using the log_2_ fold changes (LFC) values of all genes from treated versus mock samples under either DD or LD conditions at the corresponding time points with comparison groups labelled by time/light-condition/treatment (e.g., “8 DS” refers to the LFC of the samples at 8 hours under DD with SA treatment over the samples under the same conditions but with mock treatment). Pearson correlation coefficient r values (Supplementary table 2), represented in Fig. 3b by thickness of the connecting line, indicated that while SA induced similar transcriptomic changes in 8 DS and 8 LS (r = 0.76), the responses diverged substantially by 24 hours (r = 0.53 for 24 DS and 24 LS) (Fig. 3b), with a much higher number of differentially expressed genes (DEGs) at 24 DS (Extended Data Fig.3a,b), suggesting a stronger SA response under the DD condition compared to under the LD condition at 24 h. In agreement with the PCA analysis, the DSG samples showed stronger correlations with DG samples (r = 0.73 at 8 hours and 0.81 at 24 hours) than with DS samples (r = 0.49 at 8 hours and 0.6 at 24 hours), confirming that the application of glucose under DD conditions suppressed the effect of SA. However, under LD conditions, glucose had minimal impact on the SA response, as the LSG remained highly correlated with the LS samples (r = 0.79 at 8 hours and 0.82 at 24 hours), while showing weaker correlations with the LG samples (r = 0.62 at 8 hours and 0.56 at 24 hours). These differences demonstrated an emergent sensitization to SA under DD when carbohydrate supply is limited, a response that is masked during the day, in the presence of sugar produced through photosynthesis. Therefore, to eliminate the intrinsic SA-sugar crosstalk that occurs under light conditions, we focused our subsequent analyses on the interactions between SA and exogenous glucose treatments under DD, allowing for a clearer dissection of their specific contributions to and combined effects on the plant transcriptome.

**Fig. 3.**
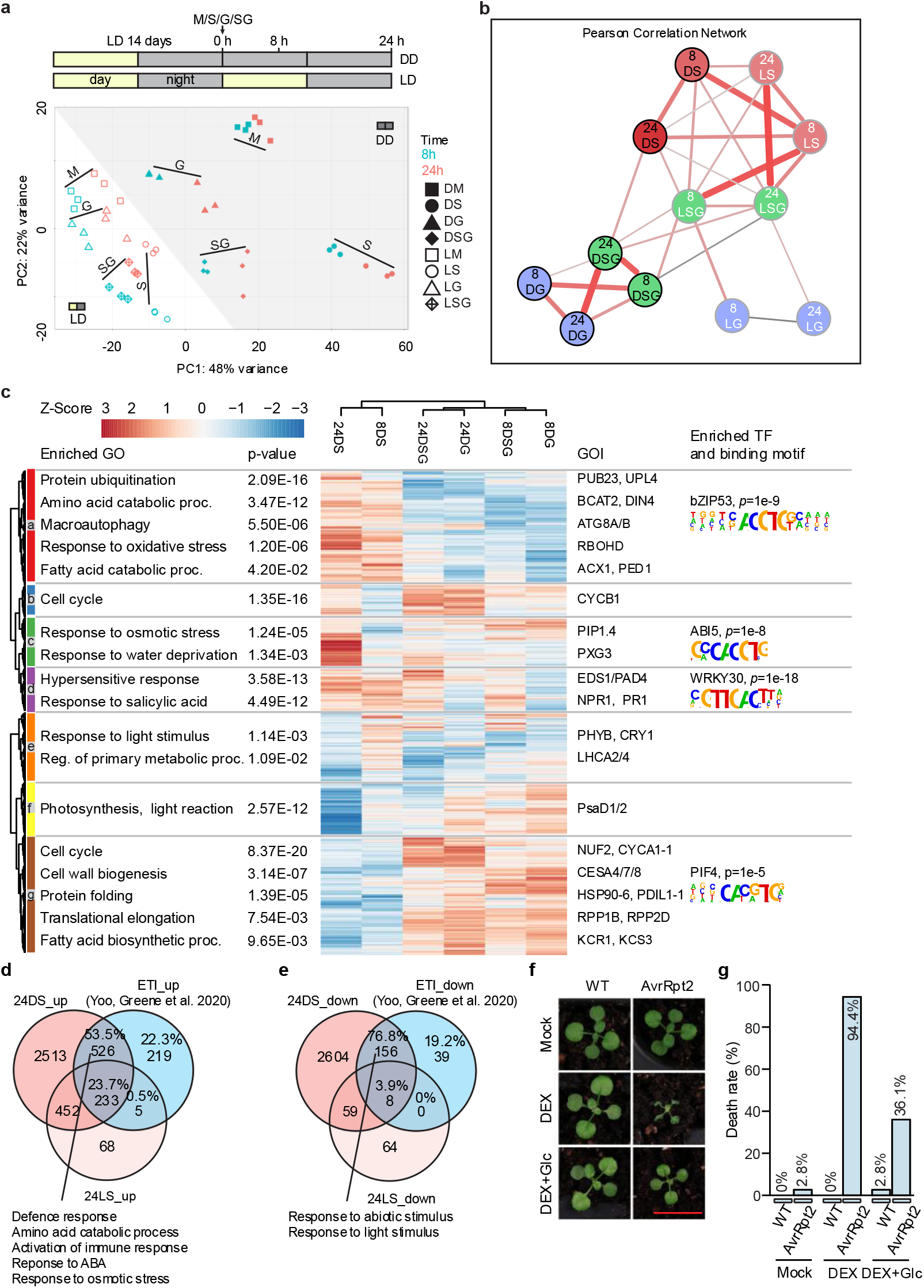
Glucose antagonizes SA-induced transcriptomic changes and rescues effector-triggered immunity (ETI)-induced plant death. (a) Schematic representation of experimental design (top) and principal component analysis (PCA) of the QuantSeq data for all the samples (bottom). Plants were grown in soil under LD conditions for 14 days, then treated with mock (M), SA (S), glucose (G) or SA plus glucose (SG) following the dark cycle. Samples were collected at 8 h and 24 h post-treatment in both LD and DD conditions. Samples in this figure are named by: timepoint (8, 24) h, light condition (D for DD; L for LD), and treatment (M, S, G, SG). (b) Pearson correlation network of log_2_ fold changes (LFC) for all samples by comparing treatment versus mock at their corresponding time points, respectively. Pearson correlation values are represented by edge colour and thickness, with a threshold of 0.5. LFC values with adjusted p-value (Padj) for all the samples are provided in Supplementary table 2. (c) Hierarchical clustering of DEGs for the DD samples. For all the figures, DEGs were defined with Padj < 0.05 and LFC > 1. The enriched regulatory motifs with their corresponding transcription factors (TFs) were generated using the HOMER v2 motif discovery software. (d and e) Venn diagrams showing the overlaps between SA up-(d) and down-(e) regulated DEGs in DD/LD conditions at 24 h with ETI up-and down-regulated genes reported previously. (f) Representative WT and dexamethasone (DEX)-inducible AvrRpt2 plants treated with mock, 0.25 µM DEX and DEX with 100 mM glucose. Plants grown in LD condition for 14 days were treated at the end of dark cycle and kept in LD for 48 h. (g) Quantification of the death rate for the plants in (f). Scale bar = 1 cm. Gene Ontology (GO) enrichment analysis was performed using both shinyGO and PANTHER GO enrichment tools with a false discovery rate (FDR) / P-value threshold of < 0.05, representative enriched terms are shown with redundant terms merged.

We first performed hierarchical clustering on all the DEGs identified by comparing treatment to mock across all conditions, using their LFC values under DD. Consistent with the Pearson correlation analysis, the DS groups clustered together, while the DSG groups clustered with the DG groups at their respective collection time points (Fig. 3c), corroborating that glucose antagonizes SA-induced transcriptomic changes. Based on their expression patterns, the DEGs were classified into seven distinct clusters (a-g). For each cluster, we carried out Gene Ontology (GO) enrichment^10,11^ followed by HOMER analysis^12^ to identify enriched biological processes and the likely regulating TFs. From these results, we highlighted selected genes of interest (GOI) and representative promoter motifs, along with their corresponding binding TFs (Fig. 3c and Supplementary table 3).

Cluster a and g represent genes up-and down-regulated by SA, respectively, whose regulation was completely reversed in the presence of glucose (Fig. 3c). GO enrichment analysis of these clusters revealed the antagonistic relationship between SA and glucose in regulating metabolism. Specifically, SA promoted catabolic process by inducing protein, amino acid, and fatty acid degradation, as well as macroautophagy, which collectively activate carbohydrate salvage pathways^13–15^. Conversely, SA supressed anabolic, growth-related processes including cell cycle, cell wall biogenesis, and protein production (translation and protein folding). Glucose reversed all these SA-mediated responses and shifted the balance from catabolism towards anabolism and growth. Promoter analysis in cluster a identified the binding sites of S1 class BASIC REGION/LEUCINE ZIPPER MOTIFs (bZIPs, e.g., bZIP53), which is known to regulate amino acid catabolism during energy deprivation^16,17^, and in cluster g the cis-elements of PHYTOCHROME INTERACTING FACTOR 4 (PIF4) ^18^, a growth promoting TF. Clusters c and f represent genes showing the strongest SA response specifically at 24 hours, which coincides with visible tissue collapse at this timepoint. These responses are also repressed by glucose, suggesting a causal relationship with plant death. The enriched terms in these clusters include increased response to osmotic stress, water deprivation and decreased photosynthesis. ABA INSENSITIVE 5 (ABI5) binding sites were overrepresented in cluster c gene promoters, consistent with their function in response to water and osmotic stress^19,20^. Cluster d included genes upregulated by SA at both 8 hours and 24 hours, while glucose had little effect on their induction. Consistent with WT-like death in the immune-deficient *npr1* mutant, this cluster is enriched with defence genes, particularly those in response to SA and ETI-triggered PCD (also known as the hypersensitive response), as well as the W-box promoter element recognized by WRKY TFs (e.g., WRKY30^21^). Clusters b and e, enriched in cell cycle and light response/primary metabolism, respectively, show expression patterns primarily driven by the time-of-day rather than treatment effects. Overall, our results showed that SA induced comparable transcriptomic changes (i.e., LFC) under DD and LD conditions at 8 h (Fig. 3b, Extended Data Fig. 3). However, at 24 hours, SA led to the accumulation of detrimental effects under DD conditions, which were attenuated by glucose treatment (Fig. 3c). Therefore, continuous glucose production from photosynthesis under LD conditions is essential for plant survival upon SA treatment.

The enrichment of genes involved ETI-induced PCD in SA-induced plant death under DD (Fig. 3c) was intriguing. To determine whether the two kinds of plant death involve common mechanisms, we compared the transcriptomic changes with the ETI data collected before the onset of PCD^22^. We first excluded those DEGs induced by SA under LD conditions (i.e., 24 LS) from the ones identified in the DD samples, because SA does not cause plant death under LD. Interestingly, we found a significant overlap between ETI-associated DEGs^22^ and DD-only SA-triggered DEGs with 53.5% and 76.8% for ETI up-and down-regulated genes, respectively (Fig. 3d,e). In addition to the expected defence gene induction, the shared upregulated gene sets were enriched in response to water or osmotic stress, and amino acid catabolic process, suggesting a positive correlation of these processes with plant death. To further test the possibility that there are shared underlying mechanisms for SA-induced death under DD and ETI-induced PCD, we applied glucose to plants after low-level dexamethasone-induced (DEX, 0.25 µM) expression of the bacterial effector AvrRpt2 and found that it could partially rescue ETI-triggered PCD (Fig. 3f,g), suggesting a critical role of sugar availability in determining SA-and ETI-induced plant death.

### The *ssd* mutants disrupt glucose metabolism and converge on downstream targets

To dissect the signalling pathway leading to the rescue of SA-induced plant death in DD by glucose, we first investigated the response in the mutants of canonical glucose sensors, HXK1 and TOR, as well as the energy sensor, SnRK1^7^. Surprisingly, we found that both the *hxk1* and the β-estradiol inducible *tor-es* silencing mutants had substantial protection against SA-induced PCD, with death rates of 26.5% and 27.8%, respectively, compared to 97.1% in the WT (Fig. 4a,b and Extended Data Fig. 4a,b), likely due to the elevated glucose and/or starch levels found in these mutants^23,24^. This result indicates that glucose rescues SA-induced plant death through a different mechanism, instead of through activities of HXK1 or TOR. Moreover, both the *snrk1.1* mutant and the *SnRK1*-overexpressing line behaved like WT (Extended Data Fig. 4c,d), ruling out the involvement of SnRK1 and the canonical energy-sensing pathway in SA-induced death.

**Fig. 4.**
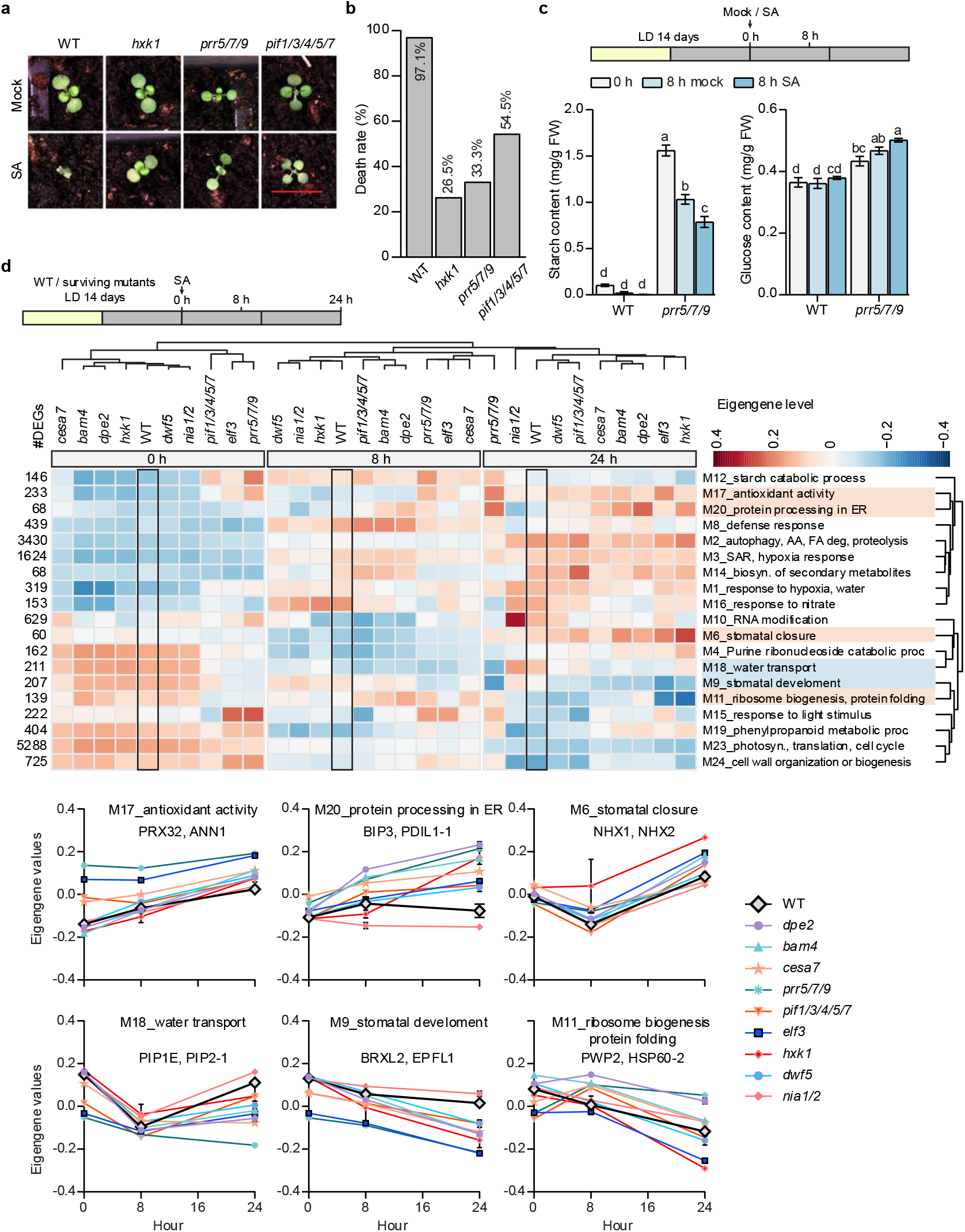
Transcriptomic analysis identifies reduced water transport and oxidative stress, and increased ER protein processing as common mechanisms for survival of SA treatment. (a) Representative images for WT, glucose sensor mutant (*hxk1*), and circadian clock-related mutants (*prr5/7/9* and *pif1/3/4/5/7*) after mock or SA treatment. Plants were grown in soil under LD conditions for 14 days, then treated with mock or SA following the dark cycle and kept in DD for 72 hours before imaging. Scale bar = 1 cm. (b) Quantification of plant death rates from (a). (c) Starch and glucose content in *prr5/7/9* mutants with mock or SA treatment in DD. The plants were grown as in (a) and were collected at 0 h and 8 h post SA treatment as shown in the schematics at the top. Six biological replicates were performed for each sample. All data are presented as mean ± s.e.m. Individual columns were compared using one-way ANOVA with Tukey’s post-hoc, different lowercase letters indicate statistical significance at p < 0.05. (d) Top: schematics of experiment design. Middle: Dendrogram combined with heatmap showed the hierarchical clustering of the eigengene values for modules identified by weighted gene co-expression network analysis (WGCNA). The heatmap was plotted using the average eigengene value for modules with a significant GO enrichment (FDR < 0.05) and without sample specificity. The DEG numbers in each module are labelled on the left and representative/top enriched GO terms are shown on the right. Columns and rows were clustered using Euclidean distance and average linkage method. Bottom: Patterns of Eigengene values for the modules displaying common differences between survival mutants and WT. Representative genes were shown for each module. AA, amino acid; FA, fatty acid.

To pinpoint the downstream signalling pathways that contribute to plant survival, we focused on the transcriptomic changes in the survival mutants. Among *ssd* mutants, the circadian clock mutant *elf3*^25^ exhibited a weak rescue phenotype (death rate of 75.8%, Fig. 1a,b), suggesting a possible functional redundancy. To investigate if the circadian clock is truly involved in the SA-induced plant death in DD, we examined the higher order clock mutant *pseudo-response regulator 5/7/9* (*prr5/7/9*)^26^, as well as the *pif1/3/4/5/7* mutant, since PIF4/5 are known circadian growth regulators^27^ downstream of ELF3. Both mutants demonstrated protection against SA-induced death under DD conditions, with *prr5/7/9* exhibiting a stronger protection (Fig. 4a,b). Importantly, we found that the *prr5/7/9* mutant had significantly higher levels of both starch and glucose compared to WT (Fig. 4c), consistent with the hypothesis that the circadian clock mutants survive due to misregulation of daily starch turnover^15^. We then performed transcriptomic analysis in representative survival mutants defective in starch and glucose metabolism (*dpe2*, *bam4*, *cesa7, hxk1*), circadian regulation (*elf3*, *prr5/7/9*, *pif1/3/4/5/7*), BR signalling (*dwf5*), and nitrate metabolism (*nia1/2*). The WT and survival mutant plants were treated with 1 mM SA at the end of the dark cycle and plants were collected at 0, 8, and 24 hours after SA treatment in DD (Fig. 4d).

PCA analyses revealed that the samples collected at the same time clustered together, indicating that all survival mutants were responsive to SA (Extended Data Fig. 4e). To identify the shared pathways that contribute to survival in all the mutants, we performed weighted gene co-expression network analysis (WGCNA) from 16,694 DEGs and, based on their expression patterns, grouped the genes into 24 co-expression modules. The eigengene values for 19 non-sample-specific modules with significant GO enrichment were then plotted as a dendrogram heatmap, together with the number of DEGs and representative GO terms (Fig. 4d and Supplementary table 4). The three most prominent modules, M2, M3 and M23, comprising the majority of the DEGs, exhibited either SA-induced (M2 and M3) or repressed (M23) expression patterns in both *ssd* mutants and WT. Since these shared modules were associated with upregulated autophagy and SAR pathways and downregulated growth-related anabolism processes, it is unlikely that growth inhibition was the direct cause of SA-induced plant death which is only observed in WT. However, in contrast to WT, all surviving mutants shared expression patterns, at at least one time point, for increased antioxidant activity, protein folding in the ER, stomata closure, and ribosome biogenesis (M17, M20, M6, and M11), along with decreased water transport and stomata development (M18 and M9), with representative genes for each module highlighted (Fig. 4d bottom). Thus, in WT, SA-induced oxidative stress and impaired water transport are associated with plant death, while in the *ssd* mutants, ribosome biogenesis and ER-protein processing serve as pro-survival mechanisms, likely by correcting oxidative stress-induced protein misfolding.

### SA induces death in DD by disrupting redox and water homeostasis

To test our hypothesis that the oxidative stress is related to SA-induced death in WT, we first measured the reduced and oxidized glutathione levels, whose ratio serves as a major redox marker reflecting cellular redox status^28^. We compared WT plants to the survival mutant *dpe2*, chosen as a representative due to its large number of *ssd* alleles, clear rescue phenotype (0% death rate), and near-WT plant size. We found that compared to the WT, SA-induced oxidation was dampened in the *dpe2* mutant, indicated by the levels of oxidized (GSSG) and reduced (GSH) glutathione as well as their ratio (GSSG/GSH) (Fig. 5a and Extended Data Fig. 5a,b). We next used H₂O₂ as an oxidation reagent, since it has been reported to induce cell death in a concentration dependent manner^29^ and SA was shown to act synergistically with H₂O₂ during ETI-triggered death PCD^30^. As expected, application of 50 mM H₂O₂, together with SA, killed 64.9% of the *dpe2* plants (Fig. 5b,c), while H_2_O_2_ alone did not cause death in either WT or the *dpe2* mutant (Extended Data Fig. 5c,d).

**Fig. 5.**
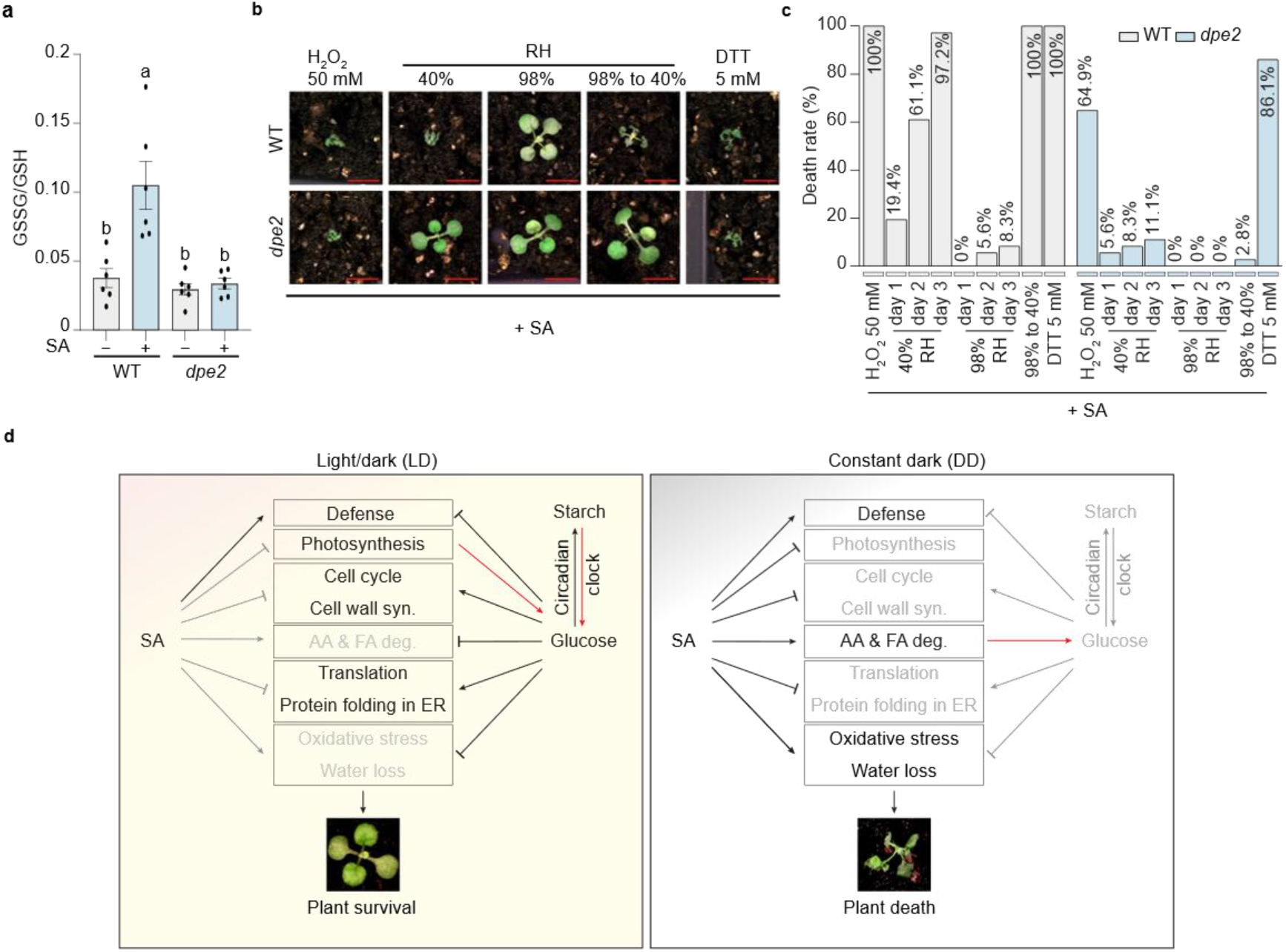
Oxidative and water stresses are the primary causes of SA-induced death. (a) GSSG/GSH ratio indicating the redox status in WT and the *dpe2* mutant. Plants grown 14 days under LD cycles were sprayed with mock or 1 mM SA at the end of dark cycle and kept in darkness. Samples were collected at 8 h post-treatment. Six biological replicates were performed for each sample. All data are presented as mean ± s.e.m. Individual columns were compared using one-way ANOVA with Tukey’s post-hoc, different lowercase letters indicate statistical significance at p < 0.05. (b) Representative images of WT and *dpe2* mutants after treatment with 1 mM SA plus 50 mM H_2_O_2_ or 5 mM DTT or only with 1 mM SA but under different relative humidity (RH) conditions. Plant grown the same way as the genetic screen were sprayed with the indicated chemicals and kept in DD for 72 h before imaging. For humidity treatment, sprayed plants were kept in a dark humidity-controlled chamber with an RH monitor. The RH was kept constant at around 40% or above 95% for 72 h before imaging. After imaging, the high RH-treated plants were moved to 40% RH under light for 4 h before images were taken once again and the death rate was calculated. Scale bar = 0.5 cm. (c) Quantification of the death rate for the plants in (b). (d) Model depicting SA-induced immune outcomes in different metabolic contexts. When WT plants are grown under LD condition, continuous carbohydrate supply from photosynthesis and starch turnover inhibits SA-induced death. Under DD, starch is depleted, SA triggers carbohydrate salvage pathways, including amino acid and fatty acid degradation, while inducing oxidative stress and water transport. The deleterious effects of SA cannot be corrected due to translation inhibition by SA and reduced ER protein folding capacity, ultimately leading to plant death. Disruption of circadian clock or primary metabolism preserves carbohydrates in DD and leads to plant survival.

Besides oxidative stress, we next tested whether preventing SA-triggered water loss would aid in WT plant survival. We found that increased relative humidity to near saturation (above 95% RH) indeed reduced WT mortality to 8.3% by day 3 in DD (Fig. 5b,c), supporting our hypothesis that increased water loss is one of the causes of SA-induced plant death in WT. This is also consistent with the enhanced death rate of *dpe2* (27.8%) by mannitol treatment compared to mock (0%), as mannitol might cause water loss through increased osmotic stress (Fig. 2c,d). Interestingly, we found that after 3 days of the high humidity treatment, returning the plants to 40% humidity for 4 hours caused 100% plant death in WT, but not in the *dpe2* mutant (Fig. 5b,c), suggesting that increased humidity blocked the water loss, perhaps the final step, in the pathway to SA-triggered plant death in DD. Similarly, ETI-triggered PCD has also been shown to be blocked by high humidity^31^ and gated by the cellular redox rhythm^28^. Together, these results demonstrate the importance of maintaining cellular redox in mitigating the metabolic toll of SA treatment and keeping water homeostasis to ensure plant survival in darkness and modulating ETI-associated PCD.

We then sought to dissect the effect of protein folding in the ER on survival. Protein folding in the ER involves two major mechanisms: disulfide bond formation, which can be blocked by DTT, and calnexin/calreticulin-mediated glycosylation, which is inhibited by tunicamycin (TM). Disrupting either pathway would trigger ER stress and activate the IRE1-dependent unfolded protein response (UPR) for protein refolding or degradation^32^. We found that, when co-sprayed with SA, TM had a marginal effect on plant fate (80.6% death in WT and 0% in *dpe2* plants), while DTT and 4μ8C (an IRE1 inhibitor) not only failed to rescue WT plants from SA-triggered death but also caused death in 86.1% and 27.8% of the *dpe2* mutants (Fig. 5b,c, and Extended Data Fig. 5e,f). When these chemicals were applied alone, without SA, little death was observed in either WT or the *dpe2* mutant. These results indicate that the disulfide bond formation pathway in protein folding in the ER is the primary pathway for promoting survival of the *ssd* mutants

Ribosome biogenesis genes were up regulated at 8 hours post SA treatment in most of the survival mutants (M11 in Fig. 4d), which might subsequently potentiate translation. In contrast, translation was downregulated in WT in response to SA treatment under DD and was rescued in response to glucose treatment (cluster g in Fig. 3c). Moreover, protein translation provides substrate as well as chaperons for protein folding in the ER^33^. These results led us to investigate whether translation itself plays a role in determining the cell fate after SA induction. Interestingly, stress caused by the translation inhibitor cycloheximide (CHX) appeared to be additive to the SA-induced stress, because CHX treatment alone was sufficient to kill WT but not *dpe2*, whereas when combined with SA, almost none of the WT or *dpe2* plants survived (Extended Data Fig. 5e,f). In contrast, inhibition of proteolysis through MG115 had little effect on SA-mediated death, indicating that proteosome does not play a major role in this response. Therefore, sustained translational activity, coupled with proper protein folding in the ER, can act as a “survival” signal to protect plants from SA-induced deleterious effects.

Finally, we tested SA analogues at 1 mM to determine whether the deleterious effects observed in this study are specific to SA. We found that application of the active analogues 2,6-dichloroisonicotinic acid (INA) killed both WT and the *dpe2* mutant in DD, while 2,1,3-benzothiadiazole (BTH), did not cause plant death even at extremely high concentration of 10 mM. This result is remarkable as BTH was formulated to minimize toxicity in plants as a commercial product^34–36^. Moreover, application of SA analogues that are inactive in triggering defence responses, 2,4-dihydroxybenzoic acid (2,4-DHBA), benzoic acid (BA), and 3-hydroxybenzoic acid (3-HBA), resulted in negligible levels of lethality (Extended Data Fig. 5g,h). The distinct effects observed between active and inactive SA analogues demonstrate that the immune response can be decoupled from the deleterious effects. To mitigate these undesirable outcomes, the pathways identified in this study can guide future investigations of the underlying signalling mechanisms.

## Discussion

SA is an important immune hormone that promotes plant survival against a broad range of pathogens and certain abiotic stresses by reprogramming physiological processes^4,37,38^. Since this involves changes in thousands of genes, how plants orchestrate SA-induced defence responses while minimizing its damage to plant growth is a central question with both fundamental and practical importance. The plant death phenotype caused by SA application in DD offers a clean and tractable experimental system to study the detrimental effects of SA which are exacerbated in the absence of light (Figs. 1 and 2f,g). The most surprising finding from this study is that defence is not the cause of death, as the *npr1* mutant, which lacks the defence responses, still perishes (Fig. 1a,b). Growth inhibition is not the cause of plant death either, because the survival mutants still showed dampened growth-related pathways, such as photosynthesis and cell cycle (e.g., M23 in Fig. 4d).

Through genetic, transcriptomic, metabolic, and chemical intervention analyses, we propose a model to explain the antagonism between SA and glucose in determining plant death or survival under different light conditions (Fig. 5d). In DD, apart from activating defence, SA induces amino acid and fatty acid degradation, water loss and oxidative stress, while simultaneously inhibiting growth-related pathways, including photosynthesis, cell cycle, cell wall biosynthesis, translation, and protein folding in the ER. Under LD conditions, continuous carbohydrate supplies from photosynthesis and starch turnover counteract SA-induced stress and support plant survival. In the survival mutants, disruption of either the circadian clock or the starch/glucose metabolism-related pathways results in increased carbohydrate levels before SA treatment, thereby preventing plant death.

The source of glucose appears to be the key determinant of plant fate. In DD, amino acids and fatty acids are mobilized to sustain glucose levels at the cost of reduced protein translation, a process identified as a hallmark of SA treatment in both darkness and LD cycle^39,40^, and shown to be associated with SA-induced death, as supported by the effect of the translation inhibitor, CHX, treatment (Extended Data Fig. 5f). Conversely, glucose derived from endogenous accumulation in the *ssd* mutants or from photosynthesis under LD is sufficient to activate the survival processes without triggering a translational or survival crisis in the plant.

The next question we asked is whether the SA-induced death under DD conditions is related to other types of plant cell death. We found that the majority of ETI-induced transcriptomic changes^22^ overlap with the SA-induced gene expression profile in DD, and that glucose can rescue plants from both SA-induced death under DD and, partially, ETI-triggered PCD in LD (Fig. 3d-g). These findings suggest that ETI-triggered PCD, leading to an energy-demanding state^41^, either results from the local increase in SA levels^42^ or from an independent mechanism, which results in death. Since in the SA-deficient mutant, ETI-triggered PCD still occurs^43^ and SA treatment alone does not normally lead to plant death under LD conditions, it is more likely that ETI involves a different induction mechanism to trigger the changes in metabolic genes. Moreover, ETI-induced PCD is also regulated by the redox metabolism^28^, understanding whether and how ETI-associated PCD is regulated in coordination with plant primary metabolic processes will be an interesting future research direction.

Through this study, it becomes more apparent how the time of day and the accompanied metabolic context determine plant fate in response to SA induction, further explaining why basal SA synthesis and plant responses to SA are directly regulated and gated by the circadian clock, respectively^1,2^. It is tempting to hypothesize that this finding may be applicable beyond plants. The SA derivative, aspirin, which is widely used in humans, such as the daily low dose of “baby aspirin” to prevent recurrent heart attacks or strokes, may exert an adverse effect if taken at an unfavourable time of the day. Elucidating the mechanisms driving the metabolic toll of SA in plants may offer valuable insights for optimizing aspirin efficacy while minimizing its potential side effects in humans where SA has been shown to also affect primary metabolic signalling^44,45^.

## Acknowledgments

X.D. would like to dedicate this study to Dr. Joanne Chory whose insightful review (<u>PMID:</u> <u>28044340</u>) “Dancing in the dark: darkness as a signal in plants” served as a source of inspiration. We thank Dr. David Somers for providing *prr5/7/9* seeds^26^, the Duke University School of Medicine for use of the Sequencing and Genomic Technologies Shared Resource for providing next-generation sequencing services and the members of the Dong laboratory for helpful discussions. We also appreciate Yetshaira Vazquez Hernandez’s assistance in the *ssd* mutant screening.

## Author contributions

Q.Z. and X.D. conceived the project. Q.Z., J.W., M.M., and H. Y. performed genetic screens and phenotypic analyses. Q.Z. and Y.X. performed transcriptomic experiments and did the data analysis. S.K. performed glutathione measurements. Q.Z. conducted chemical inhibitor and sugar intervention studies. Q.Z., Y.X., S.K. and X.D. wrote the manuscript with input from all authors.

## Data and materials availability

Requests for materials, reagents, and scripts will be addressed to the corresponding author Xinnian Dong.

## SUPPLEMENTAL INFORMATION

**Extended Data Fig. 1.**
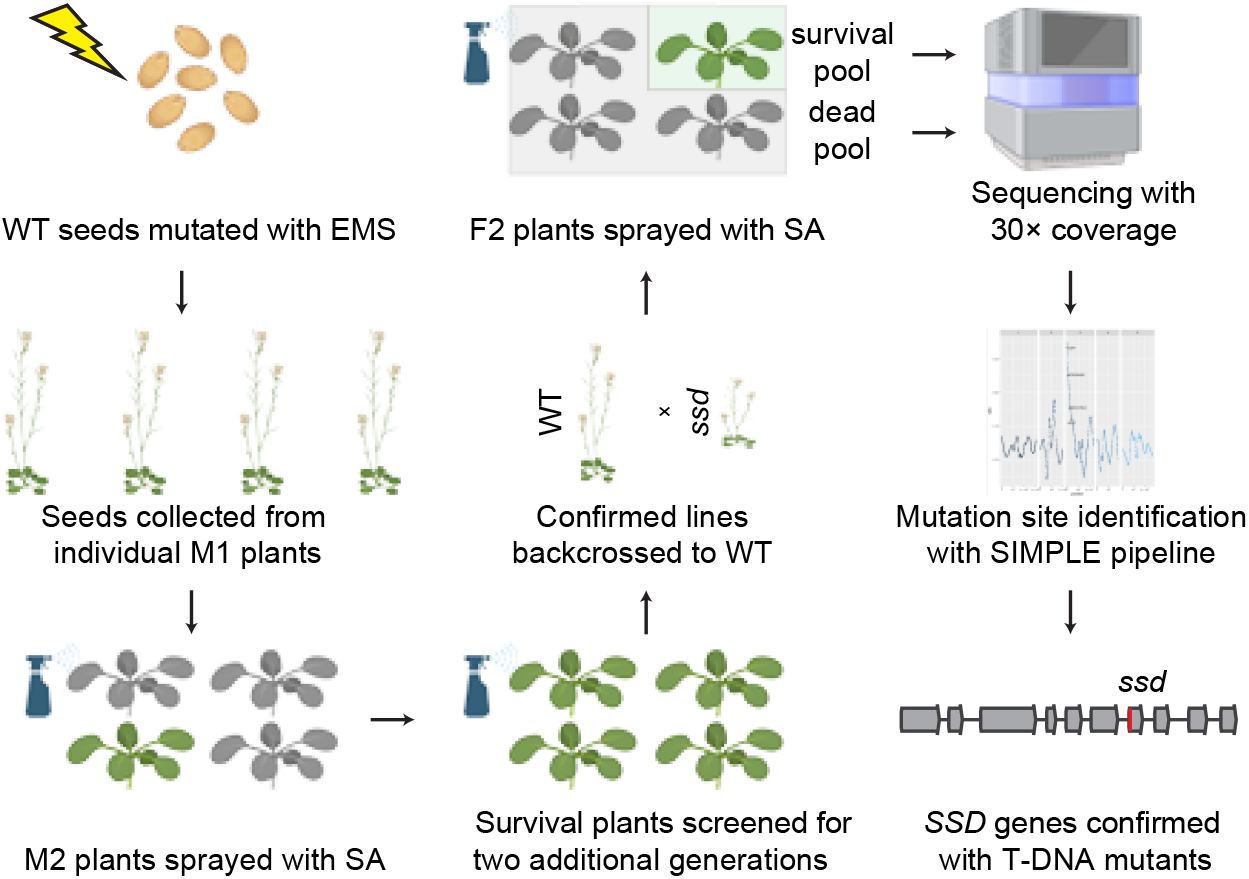
**Workflow for the genetic screen for *ssd* mutants.**WT seeds were mutagenized with EMS and sown in soil. Seeds from each M1 plant were collected separately, germinated in soil and grown for 14 days under LD cycles before treatment with 1 mM SA at the end of dark cycle. After 72 hours under DD, the surviving plants from each M2 line were collected individually and retested through two additional selfing generations. Selected surviving plants from each line were then backcrossed with WT plants to generate the F1 progeny. For each putative *ssd* line, approximately 60 surviving and 100 dead F2 plants were collected for bulk segregant whole genome sequencing at 30-fold coverage of the Arabidopsis genome. Sequencing data were analyzed using the SIMPLE pipeline to identify candidate mutations, from which the causal mutation was confirmed by examining the death rate of corresponding T-DNA insertion mutants.

**Extended Data Fig. 2.**
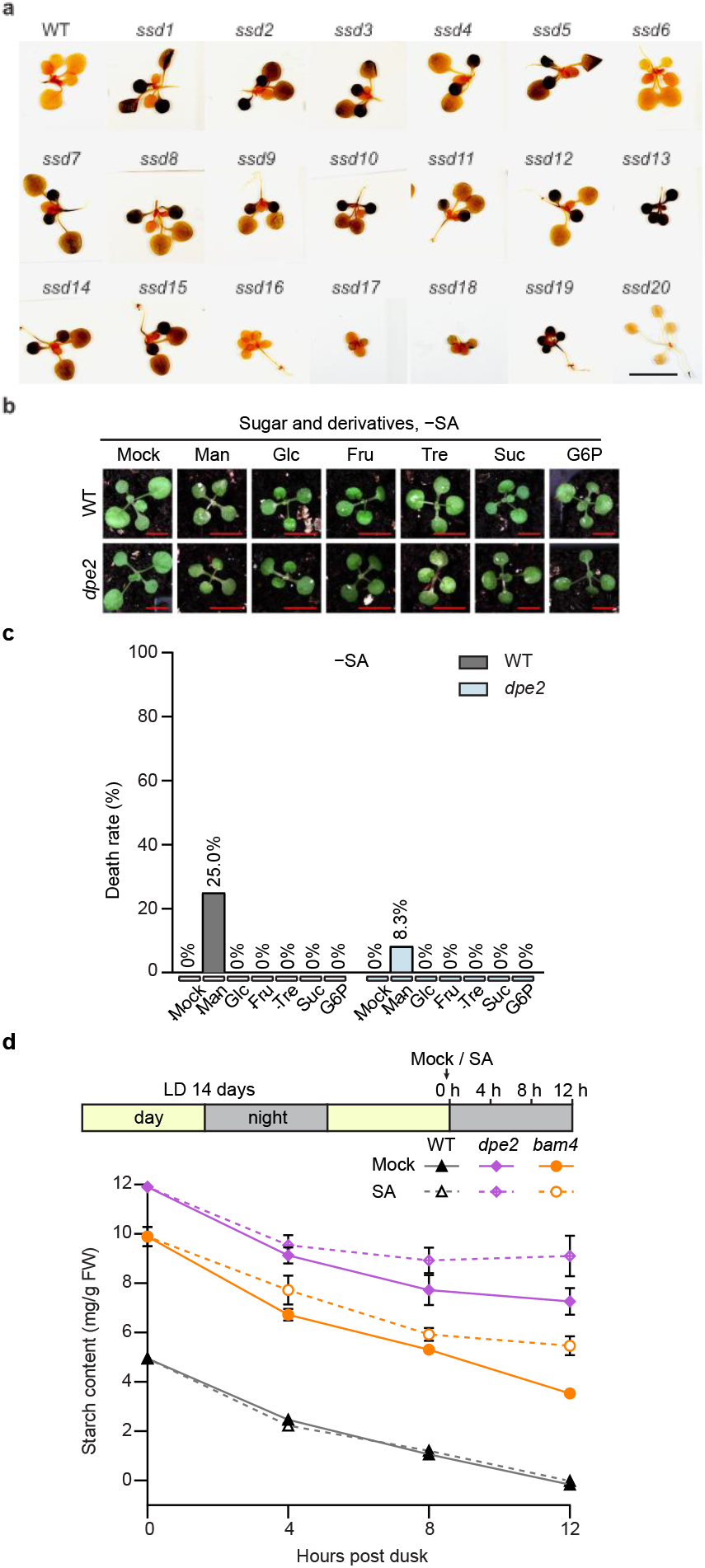
Starch measurement and SA-induced plant death rate assessments under control conditions. (a) Starch staining in WT and all the *ssd* mutants at dawn. Scale Bar = 1 cm. Plants grown for 14 days under LD cycles were collected at dawn and stained using the Lugol reagent. (b) Representative images of the plants treated with sugar and their derivatives without SA. Plants grown the same way as the genetic screen were sprayed with mock or 100 mM mannitol (Man), sugars and sugar derivatives following the dark period and kept in DD for 72 h before imaging. Glu, glucose; Fru, fructose; Tre, trehalose; G6P, glucose-6-phosphate; Scale bar = 0.5 cm. (c) Quantification of the death rate for the plants in (b). (d) Effects of SA on the starch degradation rate in WT, *dpe2*, and *bam4* mutants. Plants grown under the LD conditions for 14 days were sprayed with mock or 1 mM SA following the light cycle and samples were collected for starch quantification at 0, 4, 8, 12 h with mock or SA treatment in the dark. All data are presented as mean ± s.e.m. Six biological replicates were performed for each sample.

**Extended Data Fig. 3.**
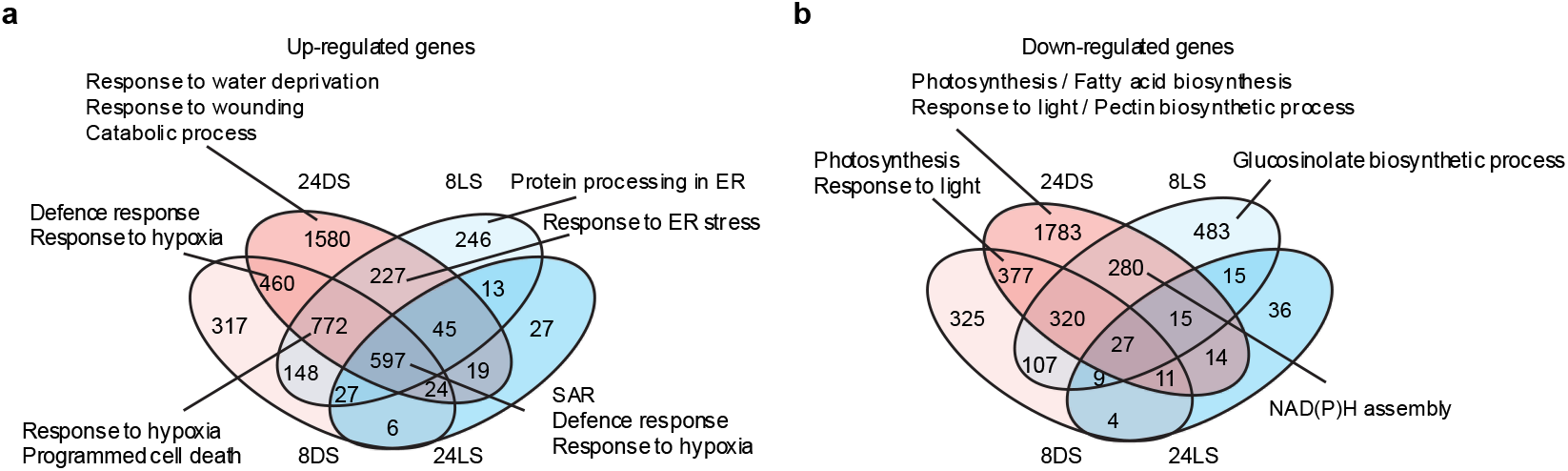
Transcriptome analysis of SA and glucose effects on plants under LD and DD conditions. (a and b) Venn diagrams showing the overlap of SA upregulated (a) and downregulated (b) DEGs between LD and DD conditions by comparing SA-to mock-treated samples at the corresponding timepoints and light conditions. Samples in this figure are designed by: timepoint (8, 24) h, light condition (D for DD; L for LD), and treatment versus mock (S, SA versus mock; G, glucose versus mock; SG, SA+glucose versus mock). DEGs were defined with Padj < 0.05 and LFC > 1

**Extended Data Fig. 4.**
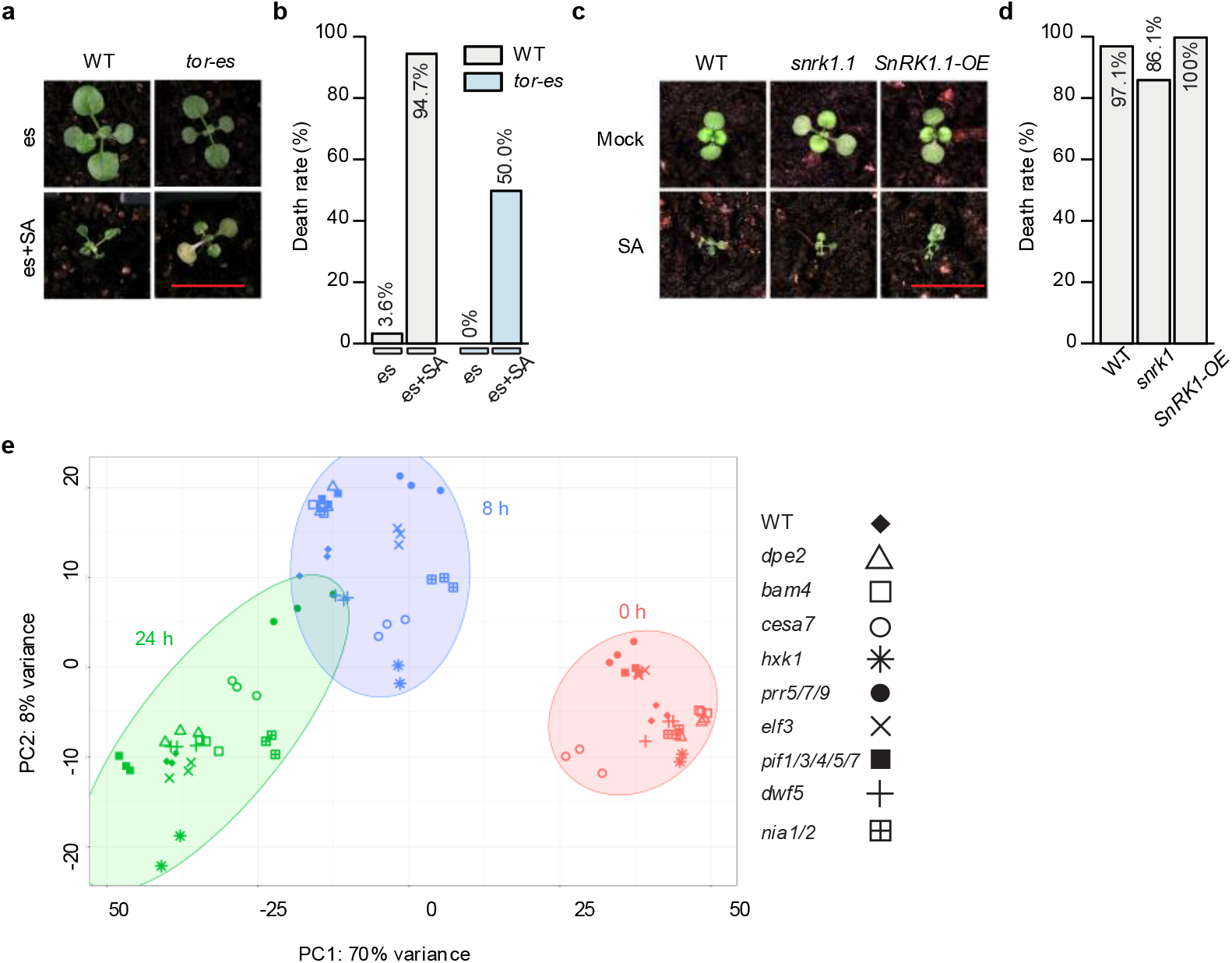
Phenotypes for sugar signalling-related mutants and PCA analysis of transcriptome data from survival mutants. (a) Representative images of WT and the *tor-es* (the β-estradiol inducible silencing line for *TOR*) plants after mock or SA treatment. Plants grown under LD cycles were pre-sprayed with 10 µM β-estradiol (es) for 5 days (one time each day) until day 14 and then treated with mock or 1 mM SA following the dark cycle. Then the plants were kept in darkness for 72 h before imaging. (b) Quantification of the death rate for the plants in (a). (c) Representative images of WT, the *snrk1.1* mutant and *SnRK1.1-OE* plants with mock or SA treatment. Plants grown for 14 days under LD cycles were sprayed with mock or 1 mM SA following the dark cycle and kept in darkness for 72 h before imaging. (d) Quantification of the death rate for the plants in (c). (e) PCA of WT and survival mutant transcriptomes with SA treatment in DD. Representative gene mutant plants were grown as in (c) and collected at 0 h, 8 h and 24 h after SA treatment in DD. Samples separated in a temporal manner in the PCA plot are marked by ellipses. Scale bar = 1 cm.

**Extended Data Fig. 5.**
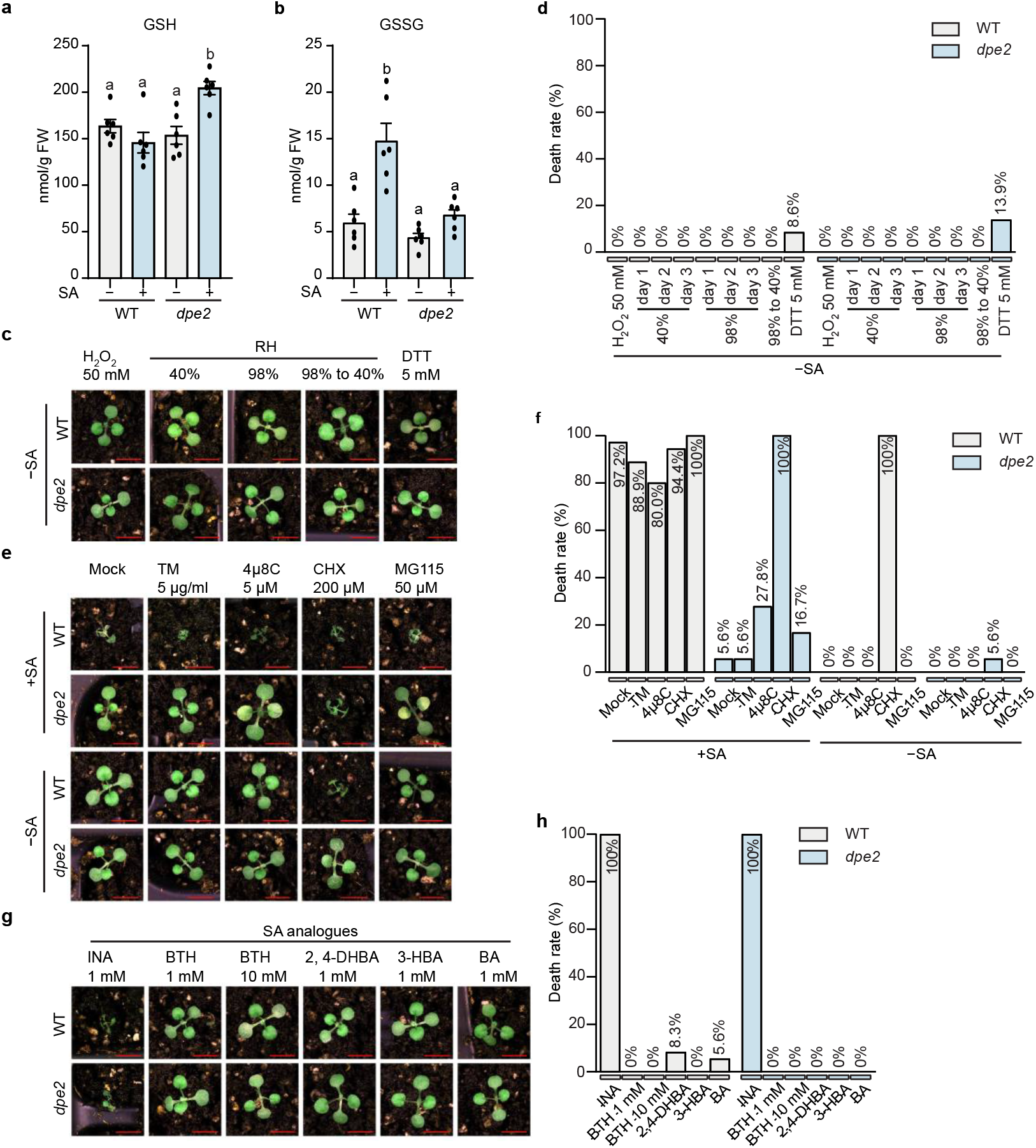
Redox measurements and additional chemical treatment phenotypes. (a and b) GSH (a), GSSG (b) measurements in WT and the *dpe2* mutant. Data are presented as means ± s.e.m. Six biological replicates were performed for each sample. Individual columns were compared using one-way ANOVA with Tukey’s post-hoc, different lowercase letters indicate statistical significance at p < 0.05. (c) Representative plant phenotypes treated with different chemicals in the absence of SA induction and under different relative humidity (RH) conditions. The treatment conditions are the same as in Figure 5B. (d) Quantification of the death rate for the plants in (c). (e) Representative plant phenotypes treated with mock or SA plus different chemicals. Plants grown for 14 days under LD cycles were treated with mock or SA plus mock, tunicamycin (TM), 4µ8C, cycloheximide (CHX) or MG115. After all treatments, plants were kept in DD for 72 h before imaging. (f) Quantification of the death rate for the plants in (e). (g) Representative plant phenotypes treated with SA active analogues [2,6-dichloropyridine-4-carboxylic acid (INA) and 2,1,3-benzothiadiazole (BTH)] or inactive analogues [2,4-dihydroxybenzoic acid (2,4-DHBA), benzoic acid (BA), and 3-hydroxybenzoic acid (3-HBA)]. Plants were grown and imaged as in (e). (h) Quantification of the death rate for the plants in (g). Scale bar = 0.5 cm.

## References

1 Zheng, X.-Y. et al. Spatial and temporal regulation of biosynthesis of the plant immune signal salicylic acid. Proceedings of the National Academy of Sciences 112, 9166–9173 (2015). 10.1073/pnas.1511182112

2 Zhou, M. et al. Redox rhythm reinforces the circadian clock to gate immune response. Nature 523, 472–476 (2015). 10.1038/nature14449

3 Powers, J. et al. Next-generation mapping of the salicylic acid signaling hub and transcriptional cascade. Molecular Plant 17, 1558–1572 (2024). 10.1016/j.molp.2024.08.008

4 Spoel, S. H. & Dong, X. Salicylic acid in plant immunity and beyond. The Plant Cell 36, 1451–1464 (2024). 10.1093/plcell/koad329

5 Li, A., Sun, X. & Liu, L. Action of Salicylic Acid on Plant Growth. Frontiers in Plant Science Volume 13–2022 (2022). 10.3389/fpls.2022.878076

6 Graf, A., Schlereth, A., Stitt, M. & Smith, A. M. Circadian control of carbohydrate availability for growth in Arabidopsis plants at night. Proceedings of the National Academy of Sciences 107, 9458–9463 (2010). doi:10.1073/pnas.0914299107

7 Sheen, J. Master Regulators in Plant Glucose Signaling Networks. J Plant Biol 57, 67–79 (2014). 10.1007/s12374-014-0902-7

8 Wachsman, G., Modliszewski, J. L., Valdes, M. & Benfey, P. N. A SIMPLE Pipeline for Mapping Point Mutations Plant Physiology 174, 1307–1313 (2017). 10.1104/pp.17.00415

9 Critchley, J. H., Zeeman, S. C., Takaha, T., Smith, A. M. & Smith, S. M. A critical role for disproportionating enzyme in starch breakdown is revealed by a knock-out mutation in Arabidopsis. The Plant Journal 26, 89–100 (2001). 10.1046/j.1365-313x.2001.01012.x

10 Ge, S. X., Jung, D. & Yao, R. ShinyGO: a graphical gene-set enrichment tool for animals and plants. Bioinformatics 36, 2628–2629 (2020). 10.1093/bioinformatics/btz931

11 Thomas, P. D. et al. PANTHER: Making genome-scale phylogenetics accessible to all. Protein Sci 31, 8–22 (2022). 10.1002/pro.4218

12 Heinz, S. et al. Simple combinations of lineage-determining transcription factors prime cis-regulatory elements required for macrophage and B cell identities. Mol Cell 38, 576–589 (2010). 10.1016/j.molcel.2010.05.004

13 Janse van Rensburg, H. C., Van den Ende, W. & Signorelli, S. Autophagy in Plants: Both a Puppet and a Puppet Master of Sugars. Front Plant Sci 10, 14 (2019). 10.3389/fpls.2019.00014

14 Yang, Q., Zhao, D. & Liu, Q. Connections Between Amino Acid Metabolisms in Plants: Lysine as an Example. Frontiers in Plant Science Volume 11–2020 (2020). 10.3389/fpls.2020.00928

15 Seluzicki, A., Burko, Y. & Chory, J. Dancing in the dark: darkness as a signal in plants. Plant Cell Environ 40, 2487–2501 (2017). 10.1111/pce.12900

16 Pedrotti, L. et al. Snf1-RELATED KINASE1-Controlled C/S(1)-bZIP Signaling Activates Alternative Mitochondrial Metabolic Pathways to Ensure Plant Survival in Extended Darkness. Plant Cell 30, 495–509 (2018). 10.1105/tpc.17.00414

17 Garg, A. et al. Targeted manipulation of bZIP53 DNA-binding properties influences Arabidopsis metabolism and growth. Journal of Experimental Botany 70, 5659–5671 (2019). 10.1093/jxb/erz309

18 Sun, J., Qi, L., Li, Y., Chu, J. & Li, C. PIF4–Mediated Activation of YUCCA8 Expression Integrates Temperature into the Auxin Pathway in Regulating Arabidopsis Hypocotyl Growth. PLOS Genetics 8, e1002594 (2012). 10.1371/journal.pgen.1002594

19 Ramegowda, V. et al. GBF3 transcription factor imparts drought tolerance in Arabidopsis thaliana. Scientific Reports 7, 9148 (2017). 10.1038/s41598-017-09542-1

20 Skubacz, A., Daszkowska-Golec, A. & Szarejko, I. The Role and Regulation of ABI5 (ABA-Insensitive 5) in Plant Development, Abiotic Stress Responses and Phytohormone Crosstalk. Frontiers in Plant Science Volume 7–2016 (2016). 10.3389/fpls.2016.01884

21 Zou, L. et al. Transcription factor WRKY30 mediates resistance to Cucumber mosaic virus in Arabidopsis. Biochemical and Biophysical Research Communications 517, 118–124 (2019). 10.1016/j.bbrc.2019.07.030

22 Yoo, H. et al. Translational Regulation of Metabolic Dynamics during Effector-Triggered Immunity. Molecular Plant 13, 88–98 (2020). 10.1016/j.molp.2019.09.009

23 Brauner, K., Stutz, S., Paul, M. & Heyer, A. G. Measuring whole plant CO2 exchange with the environment reveals opposing effects of the gin2-1 mutation in shoots and roots of Arabidopsis thaliana. Plant Signal Behav 10, e973822 (2015). 10.4161/15592324.2014.973822

24 Caldana, C. et al. Systemic analysis of inducible target of rapamycin mutants reveal a general metabolic switch controlling growth in Arabidopsis thaliana. Plant J 73, 897–909 (2013). 10.1111/tpj.12080

25 Liu, X. L., Covington, M. F., Fankhauser, C., Chory, J. & Wagner, D. R. ELF3 encodes a circadian clock-regulated nuclear protein that functions in an Arabidopsis PHYB signal transduction pathway. Plant Cell 13, 1293–1304 (2001). 10.1105/tpc.13.6.1293

26 Wang, L., Kim, J. & Somers, D. E. Transcriptional corepressor TOPLESS complexes with pseudoresponse regulator proteins and histone deacetylases to regulate circadian transcription. Proceedings of the National Academy of Sciences 110, 761–766 (2013). doi:10.1073/pnas.1215010110

27 Box, Mathew S. et al. ELF3 Controls Thermoresponsive Growth in Arabidopsis. Current Biology 25, 194–199 (2015). 10.1016/j.cub.2014.10.076

28 Karapetyan, S., Mwimba, M., Chen, T., Yao, Z. & Dong, X. The redox rhythm gates immune-induced cell death distinctly from the genetic clock. Proceedings of the National Academy of Sciences 122, e2519251122 (2025). doi:10.1073/pnas.2519251122

29 Desikan, R., Reynolds, A., Hancock, J. T. & Neill, S. J. Harpin and hydrogen peroxide both initiate programmed cell death but have differential effects on defence gene expression in Arabidopsis suspension cultures. Biochem J 330 ( Pt 1), 115–120 (1998). 10.1042/bj3300115

30 Shirasu, K., Nakajima, H., Rajasekhar, V. K., Dixon, R. A. & Lamb, C. Salicylic acid potentiates an agonist-dependent gain control that amplifies pathogen signals in the activation of defense mechanisms. Plant Cell 9, 261–270 (1997). 10.1105/tpc.9.2.261

31 Xin, X.-F. et al. Bacteria establish an aqueous living space in plants crucial for virulence. Nature 539, 524–529 (2016). 10.1038/nature20166

32 Howell, S. H. Endoplasmic reticulum stress responses in plants. Annu Rev Plant Biol 64, 477–499 (2013). 10.1146/annurev-arplant-050312-120053

33 Dabsan, S., Twito, G., Biadsy, S. & Igbaria, A. Less is better: various means to reduce protein load in the endoplasmic reticulum. The FEBS Journal 292, 976–989 (2025). 10.1111/febs.17201

34 Friedrich, L. et al. A benzothiadiazole derivative induces systemic acquired resistance in tobacco. The Plant Journal 10, 61–70 (1996). 10.1046/j.1365-313X.1996.10010061.x

35 Görlach, J. et al. Benzothiadiazole, a novel class of inducers of systemic acquired resistance, activates gene expression and disease resistance in wheat. The Plant Cell 8, 629–643 (1996). 10.1105/tpc.8.4.629

36 Lawton, K. A. et al. Benzothiadiazole induces disease resistance in Arabidopsis by activation of the systemic acquired resistance signal transduction pathway. The Plant Journal 10, 71–82 (1996). 10.1046/j.1365-313X.1996.10010071.x

37 Liu, J. et al. Salicylic Acid, a Multifaceted Hormone, Combats Abiotic Stresses in Plants. Life (Basel) 12 (2022). 10.3390/life12060886

38 Zavaliev, R. & Dong, X. NPR1, a key immune regulator for plant survival under biotic and abiotic stresses. Mol Cell 84, 131–141 (2024). 10.1016/j.molcel.2023.11.018

39 Chen, Y. et al. Salicylic Acid Engages Central Metabolic Regulators SnRK1 and TOR to Govern Immunity by Differential Phosphorylation of NPR1. bioRxiv, 2025.2006.2017.660129 (2025). 10.1101/2025.06.17.660129

40 Xie, Z. et al. Phenolic acid-induced phase separation and translation inhibition mediate plant interspecific competition. Nat Plants 9, 1481–1499 (2023). 10.1038/s41477-023-01499-6

41 Chen, T. et al. Global translational induction during NLR-mediated immunity in plants is dynamically regulated by CDC123, an ATP-sensitive protein. Cell Host Microbe 31, 334–342.e335 (2023). 10.1016/j.chom.2023.01.014

42 Betsuyaku, S. et al. Salicylic Acid and Jasmonic Acid Pathways are Activated in Spatially Different Domains Around the Infection Site During Effector-Triggered Immunity in Arabidopsis thaliana. Plant Cell Physiol 59, 8–16 (2018). 10.1093/pcp/pcx181

43 Cui, H. et al. A core function of EDS1 with PAD4 is to protect the salicylic acid defense sector in Arabidopsis immunity. New Phytologist 213, 1802–1817 (2017). 10.1111/nph.14302

44 Din, F. V. et al. Aspirin inhibits mTOR signaling, activates AMP-activated protein kinase, and induces autophagy in colorectal cancer cells. Gastroenterology 142, 1504–1515.e1503 (2012). 10.1053/j.gastro.2012.02.050

45 Hawley, S. A. et al. The ancient drug salicylate directly activates AMP-activated protein kinase. Science 336, 918–922 (2012). 10.1126/science.1215327

